# V genes in primates from whole genome shotgun data

**DOI:** 10.1101/006924

**Authors:** David N. Olivieri, Francisco Gambón-Deza

## Abstract

The adaptive immune system uses V genes for antigen recognition. The evolutionary diversification and selection processes within and across species and orders are poorly understood. Here, we studied the amino acid (AA) sequences obtained of translated in-frame V exons of immunoglobulins (IG) and T cell receptors (TR) from 16 primate species whose genomes have been sequenced. Multi-species comparative analysis supports the hypothesis that V genes in the IG loci undergo birth/death processes, thereby permitting rapid adaptability over evolutionary time. We also show that multiple cladistic groupings exist in the TRA (35 clades) and TRB (25 clades) V gene loci and that each primate species typically contributes at least one V gene to each of these clade. The results demonstrate that IG V genes and TR V genes have quite different evolutionary pathways; multiple duplications can explain the IG loci results, while co-evolutionary pressures can explain the phylogenetic results, as seen in genes of the TR loci. We describe how each of the 35 V genes clades of the TRA locus and 25 clades of the TRB locus must have specific and necessary roles for the viability of the species.

## 1 Introduction

The adaptive immune system contains natural molecular recognition machinery that is able to distinguish self from non-self and defend the body against infections (Janeway Jr, 1992). This molecular recognition system consists of two molecular structures, immunoglobulins (IG) and T lymphocyte receptors (TR). Immunoglobulins recognize antigen in soluble form and are composed of two types of molecular units, a heavy chain (IGH) and a light chain (either IGK or IGL). The recognition site is composed of the variable (V) domains present in the NH2-terminus of both chains. The antigen binding site is composed of two V domains, one from each chain. Within these protein domains, zones have been described that interact with antigen (called the complementarity determining regions, CDR) and framework regions (FR). For IG, there are three CDR and three FR regions within each V domain. The interaction site with antigen consists of six CDR supported by six FR regions. The TR recognize antigen that are presented by the molecules of the major histocompatibility complex (MHC), as antigen-MHC molecular complexes. Despite this substantial difference between TR and IG with respect to the mechanism of antigen recognition, both possess similar structures. In particular, each have two chains possessing V domains at the site within the molecule that is responsible for antigen-MHC recognition. These domains are similar to those in IG, containing three FR regions and three CDR per chain (Janeway et al., 2001).

Because the amino acid (AA) sequences of the IG and TR V domains are so similar, it has been hypothesized that all such sequences present today were derived from an ancestral gene (Hughes, 1994). This ancestral gene was assigned to immune recognition in the epoch coinciding with the origin of vertebrates. Evidence comes from the present-day IG and TR sequences in fish, whose structures have been maintained in all extant vertebrates (Ghaffari & Lobb, 1991).

Genes of the V domains are distributed across seven unique loci. The genes from three of these loci are used to construct IG chains, while the other four loci contain genes that encode TR chains (Janeway et al., 2005; Brack et al., 1978; Tonegawa, 1983; Davis & Bjorkman, 1988). The antigen recognition repertoire that an organism possesses is dictated by the total available set of these genes. These genes have two exons, one for the peptide leader and the other that encodes most of the V domain (V in-frame exon) (The Immunoglobulin FactsBook (Lefranc & Lefranc, 2001a); The T cell receptor FactsBook (Lefranc & Lefranc, 2001b); (Lefranc, 2014). Within the V exon, there are coding sequences for the first two complementarity determining region (CDR1 and CDR2) for antigen recognition and the three framework regions (FR1, FR2, and FR3) (the international ImMunoGeneTics information system, http://www.imgt.org (Lefranc et al., 2009), IMGT/GENE-DB (Giudicelli & Lefranc, 2004). A third complementarity determining region (CDR3) is generated through a gene rearrangement process, called VDJ recombination, whereby V exons are moved from their location in order to join with other gene segments, called D and J. This process is somatic and only occurs within lymphoid cells (Tonegawa, 1983).

The number of V exons for IG and TR is highly variable across different species, especially with respect to the IG loci. For example, there are approximately 600 IGHV genes for the microbat (*Mioitis lucifugus*), while for other mammals, such as those living in aquatic environments (e.g. seals, dolphins and walruses) have much fewer IGHV genes (Olivieri et al., 2013). From the V exon sequence data available at http://vgenerepertoire.org, the number of genes in the TRBV locus between species is approximately constant, while there is a large variance in the number of TRAV genes, particularly pronounced in the Bovine species of the Laurasiatheria. The causes for this variability amongst species is presently unknown.

In this paper, we describe organizational and phylogenetic relationships of the amino acid (AA) sequences derived from the V exons of the order Primates uncovered from whole genome shotgun (WGS) datasets. In particular, we studied 16 representative Primate species whose genomes have been sequenced in order to identify evolutionary patterns that could explain the present-day genomic repertoire of V genes. Our results show that in the IG loci, duplications and losses of V exons are common, while in the TRAV and TRBV loci, complex selection mechanisms may be responsible in order to conserve V exons between species.

## 2 Material y methods

For these studies, we used genome data from whole genome shotgun (WGS) assemblies of species that are publicly available at the NCBI. For the majority of these species, the V genes have not been annotated or only partial annotations have been performed in specific loci, (IMGT Repertoire, http://www.imgt.org). All curated genes were entered in IMGT/GENE-DB (Giudicelli et al., 2005) and IMGT gene nomenclature has been provided to Gene at NCBI (Lefranc, 2014). We used our VgenExtractor bioinformatics tool [http://vgenerepertoire.org/downloads/] (Olivieri et al., 2013) to identify the in-frame V exon sequences from an analysis of these genome files by searching for well established signatures and motifs. Our software algorithm searches and extracts in-frame V exon sequences based upon known motifs. In particular, the algorithm scans large genome files and extracts candidate V exon sequences. These V exon sequences are delimited at the nucleotide level by an acceptor splice a the 5′ end and at the 3′ end by the V recombination signal (V-RS). These V exon (and its translation) includes the second part of the signal peptide (L-PART2) and the V-REGION (IMGT-ONTOLOGY 2012, for a detailed description (Giudicelli & Lefranc, 2012)). The IMGT unique numbering starts at the beginning of the V-REGION. The exons fulfill specific criteria: they are flanked in 3′ by the V recombination signal (V-RS), they have a reading frame without stop codon, the length is at least 280 bp long, and they contain two canonical cysteines and a tryptophan at specific positions (1st-CYS 23 and 2nd-CYS 104, CONSERVED-TRP 41 according to the IMGT unique numbering ((Lefranc et al., 2003), (Lefranc, 2011)).

Since the VgenExtractor algorithm scans entire genomes by matching specific motif patterns along the exon sequence, a fraction of the extracted sequences can fulfill the conditions of our algorithm for being functional V-genes, yet are structurally very different. Such sequences are easily discarded with a Blastp comparison against a V-gene consensus sequence. We found that an ample threshold (evalue=1e-15) is sufficient for eliminating all sequences that are not V genes.

The VgenExtractor algorithm can be modified to identify pseudogenes by relaxing the motif filters or by relaxing the condition of stop-codon translation. Nonetheless, this would only uncover a fraction of the complete set of pseudogenes that could otherwise fulfill different criteria. A complete set of pseudogenes would remain elusive due to random alterations of sequences over evolutionary history. Thus, we limited all further study to specifically targeted functional genes, or those exons that possess the requirements seen in all V genes sequences annotated to date (IMGT Repertoire, http://www.imgt.org).

Once we identified functional V exons, we analyzed the set of translated amino acid (AA) sequences with a pipeline that we implemented within the Galaxy toolset (https://usegalaxy.org/). The steps of the workflow are as follows. First, we performed multiple BLAST alignment of the AA sequences against V exon consensus sequences obtained from previously annotated V genes (IMGT Repertoire, http://www.imgt.org). Those sequences with a BLAST similarity score > 0.001 were retained, while other sequences were discarded. From the resulting AA translated V exon sequences, we performed multiple alignment with ClustalO(Sievers & Higgins, 2014) and phylogenetic comparison studies using SEAVIEW (Gouy et al., 2010). For the tree construction, we used a maximum likelihood algorithm and the LG matrix. Finally, we used the MEGA5 (Tamura et al., 2011) and FigTree (http://tree.bio.ed.ac.uk/software/figtree/) to produce tree graphics.

We classified the V exon sequences into one of the 7 loci (IGHV, IGKV, IGLV, TRAV, TRBV, TRDV and TRGV) by obtaining a heuristic score based upon a BLASTP against the NCBI NR protein database. The score is computed by mining the text description from protein hits that have a similarity score above a predetermined threshold in order to obtain a relevant word frequency indicative of exon type. For each protein description, the word frequency is weighted by the BLAST similarity score, so that the most similar protein descriptions contribute more to the final loci classification.

We developed a python analysis script (called *Trozos*, which can be freely downloaded at http://vgenextractor.org/downloads/), which we used for extracting the CDR and FR sequences from the AA translated V exon. First, we studied the V exon amino acid sequences and could identify the presence of the two canonical cysteines and the presence of tryptophan, W41 (IMGT unique numbering (Lefranc et al., 2003; Lefranc, 2011)). Between each V exon, the number of amino acids in the CDRs varies, however we used the standardized IMGT naming/nomenclature to define the regions. In particular, CDR1 contains six amino acids and begins at the position of the first cysteine (ie., + 3 to cysteine + 10). The CDR2 is defined as the sequence located between W41 + 15 and W41 + 22. The framework sequence regions are located between the CDRs. Stretches of sequences, which we indicate by (i), refer to sequences we obtained computationally. For a particular set of sequences, some parameter adjustment in the *Trozos* algorithm is necessary for consistency, validating the final result against a visualization of the sequence alignment. For a detailed study of the alignments and a study of the conservation sites, we used the software Jalview (Waterhouse et al., 2009).

We obtained the V exon sequences of 16 primates whose WGS sequences are available at the NCBI. The primates included in our study and their corresponding abbreviated accession numbers are the following: the Lemuriformes: *Daubentonia madagascariensis* (AGTM01), *Otolemur garnettii* (AAQR03), *Microcebus murinus* (AAHY01), the Tarsiformes: *Tarsius syrichta* (ABRT01), the New World Monkeys: *Callithrix jacchus* (ACFV01), *Saimiri boliviensis* (AGCE01), the Old World Monkeys: *Macaca mulatta* (AANU01), *Macaca fascicularis* (CAEC01), *Chlorocebus sabaeus* (AQIB01), *Papio anubis* (AHZZ01), the Hominids: *Nomascus leucogenys* (ADFV01), *Pongo abelii* (ABGA01), *Gorilla gorilla* (CABD02), *Pan paniscus* (AJFE01), *Pan troglodytes* (AACZ03), and *Homo sapiens* (AADD01). Details of these WGS data sets are provided in Supplementary Table 1.

**Table 1:** Distribution of V-genes amongst the IG and TR loci.

| SPECIE | IGHV | IGKV | IGLV | TRAV | TRBV | TRGV | TRDV | ALL |
| --- | --- | --- | --- | --- | --- | --- | --- | --- |
| LEMURIFORMES |  |  |  |  |  |  |  |  |
| *D. madagascariensis* | 4 | 10 | 10 | 40 | 17 | 2 | 5 | 88 |
| *O. garnettii* | 9 | 49 | 42 | 69 | 31 | 5 | 13 | 218 |
| *M. murinus* | 40 | 26 | 52 | 38 | 30 | 3 | 3 | 191 |
| TARSIIFORMES |  |  |  |  |  |  |  |  |
| *T. syrichta* | 38 | 50 | 26 | 40 | 12 | 3 | 6 | 175 |
| NEW WORLD MONKEYS |  |  |  |  |  |  |  |  |
| *C. jacchus* | 75 | 24 | 27 | 30 | 43 | 3 | 6 | 208 |
| *S. boliviensis* | 27 | 29 | 14 | 37 | 36 | 4 | 2 | 159 |
| OLD WORLD MONKEYS |  |  |  |  |  |  |  |  |
| *M. mulatta* | 63 | 65 | 51 | 45 | 57 | 5 | 6 | 292 |
| *M. fascicularis* | 67 | 66 | 48 | 44 | 56 | 4 | 7 | 292 |
| *C. sabaues* | 64 | 63 | 44 | 36 | 53 | 3 | 7 | 270 |
| *P. anubis* | 68 | 78 | 47 | 38 | 58 | 5 | 6 | 300 |
| HOMINIDS |  |  |  |  |  |  |  |  |
| *N. leucogenys* | 28 | 33 | 20 | 42 | 37 | 4 | 5 | 169 |
| *P. abelii* | 72 | 42 | 31 | 42 | 62 | 4 | 9 | 262 |
| *G. gorilla* | 44 | 30 | 22 | 41 | 49 | 3 | 5 | 194 |
| *P. paniscus* | 30 | 26 | 28 | 36 | 51 | 6 | 6 | 183 |
| *P. troglodytes* | 53 | 41 | 27 | 52 | 55 | 5 | 4 | 237 |
| *H. sapiens* | 44 | 46 | 33 | 42 | 49 | 3 | 6 | 193 |

## 3 Results

Previous work have described variations in the number of V genes between species (Guo et al., 2011; Niku et al., 2012). Likewise, we recently demonstrated the presence of distinct evolutionary processes between the IG and TR V genes (Olivieri et al., 2013). Nonetheless, the origin of this variation is still unknown. In order to understand the reason for this V gene number variation, we compared these genes amongst species of specific mammalian orders and families. Here we describe such a comparative study in primates. In particular, we studied the V exon sequences from the 16 primate species represented in Figure 1. We extracted the V exon sequences from WGS public datasets, listed in Table 1. We carried out studies in the five major simian branches. Nonetheless, there is a greater representation (six species) from the hominid group simply based upon the maturity of available WGS data.

**Figure 1:**
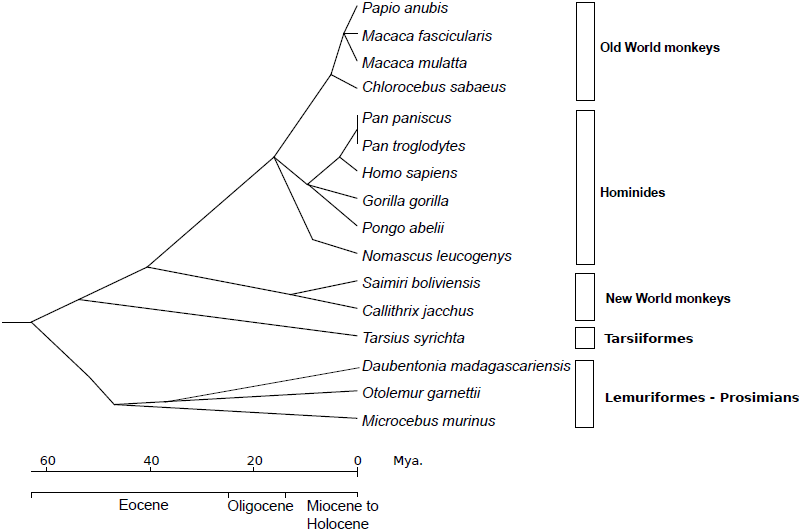
Phylogenetic tree of the primates included in the study. The tree was constructed from recent molecular phylogenetic data provided in (Perelman et al., 2011).

### 3.1 Immunoglobulin Genes

The IGHV locus has been described in many vertebrate species (Berman et al., 1988; Miller et al., 1998; Deza et al., 2009; Gambón-Deza et al., 2010). The joint study of all species with all V exon sequences has identified three evolutionary clans (IMGT (Lefranc, 2001)). The clans are defined by specific sequences in the Framework 1 (IMGT-FR1) regions and Framework 3 (IMGT-FR3) regions, which are influential in antigen recognition functionality (Kirkham et al., 1992). The IGKV and IGKL loci are less well studied and there are no published work that clearly indicate the existence of clans in these loci, as is the case in the IGHV locus.

From the 16 species of primates studied (Table 1), we obtained a total of 701 IGHV exons sequences. Two species, (*D. madagascariensis* and *O. garnetti*), have markedly fewer IGHV genes than the average, while the rest of the primates have an average of between 30 to 60 IGHV genes in this locus.

To compare the V exon sequences in extant primate species, we carried out multi-species phylogenetic analysis by first aligning the AA translated sequences with clustalO (Sievers & Higgins, 2014) and then performing tree construction with FastTree (Price et al., 2010) (using maximum likelihood and WAG matrices). Subsequently, we used Figtree or MEGA5 (Tamura et al., 2011) for visualization. The resulting phylogenetic trees show the presence of three major clades (Clan I, II and III) which have already been described in vertebrates (Kirkham et al., 1992; Lefranc & Lefranc, 2001a; Giudicelli & Lefranc, 1999) (see the IMGT clans http://www.imgt.org) (Figure 2). Additionally, due to the large number of sequences in this study, we can discern the presence of subclades within each of the defined IGHV locus clans. Thus, from the primate V exon sequences found within Clan I, the 182 sequences form three subclades (A-29 Seq, B-22 Seq, and C-131 Seq), the 139 sequences in Clan II form two subclades (A-41 Seq and B-98 Seq), and the 380 sequence in Clan III 380 can be grouped into three subclades (A-104 Seq, B-94 Seq, and C-182 Seq).

**Figure 2:**
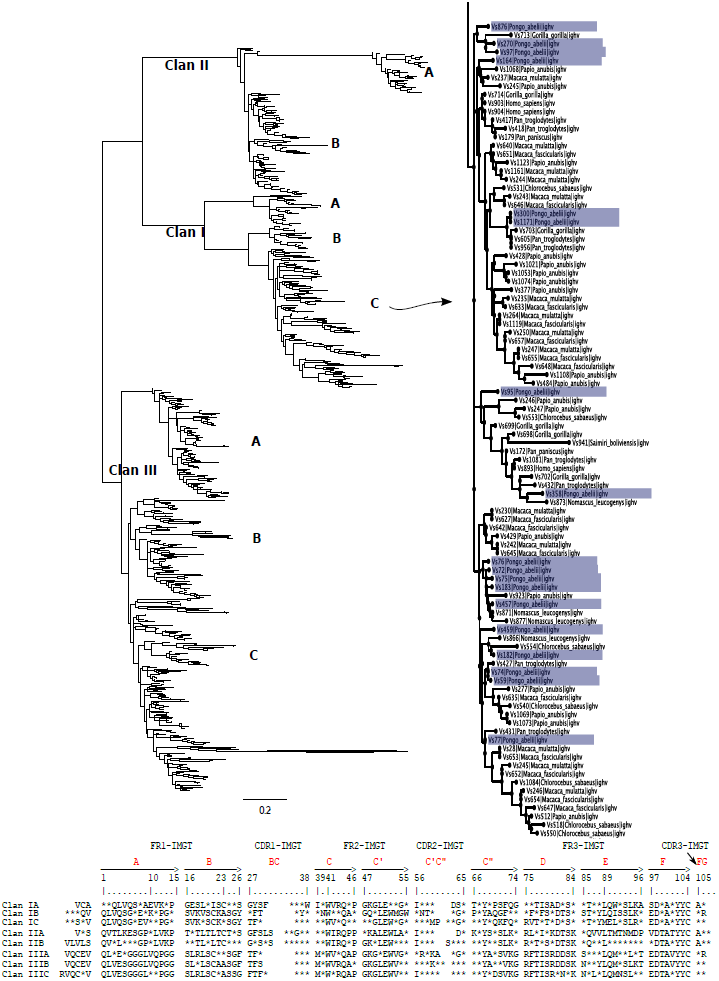
The phylogenetic trees of the AA translated sequences from IGHV exons from 16 primates species. V exon sequences are obtained from whole genome shotgun (WGS) datasets using the Vgenextractor algorithm. Alignment of the amino acid sequences was performed with clustalO, tree construction with FastTree using the WAG matrix, and visualization with Figtree. (Left:) the tree of all IGHV exon sequences; (Right:) a detailed view of the subclade, Clan I-C, showing the names of constituent species. The leaves of the taxon, *P. albelii*, are highlighted in order to easily illustrate the distribution of a particular species within the subclade. In the bottom part of the figure, the consensus sequences of each clade are given. The amino acids that are found in more than 90 % of the sequences are marked by their letter, while the variable regions are represented by an asterisk (“*”).

Clues about the origins of V gene sequences can be gained by observing the distribution of the primates amongst the clades. While Old World monkeys and Hominids have sequences in all clades, New World Monkeys have no sequences within clade III-B and Tassiforms and Lemures have no sequences within the IGHV clade II (Table 2). We also observe that species typically have several V genes per clade, as seen in Figure 2), which may be due to recent duplication events.

**Table 2:** Distribution of IGHV exons across the clans.

|  | CLAN I |  |  | CLAN II |  | CLAN III |  |  |
| --- | --- | --- | --- | --- | --- | --- | --- | --- |
| SPECIE | A | B | C | A | B | A | B | C |
| LEMURIFORMES |  |  |  |  |  |  |  |  |
| *D. madagasca.* | 0 | 0 | 2 | 0 | 0 | 0 | 0 | 0 |
| *O. garnettii* | 2 | 0 | 0 | 0 | 0 | 2 | 4 | 1 |
| *M. murinus* | 0 | 0 | 9 | 0 | 0 | 5 | 10 | 13 |
| TARSIIFORMES |  |  |  |  |  |  |  |  |
| *T. syrichta* | 2 | 0 | 16 | 0 | 0 | 5 | 8 | 8 |
| NEW WORLD MONKEYS |  |  |  |  |  |  |  |  |
| *C. jacchus* | 7 | 2 | 18 | 1 | 2 | 3 | 0 | 40 |
| *S. boliviensis* | 2 | 2 | 8 | 2 | 1 | 2 | 0 | 10 |
| OLD WORLD MONKEYS |  |  |  |  |  |  |  |  |
| *M. mulatta* | 2 | 1 | 4 | 4 | 14 | 14 | 9 | 15 |
| *M. fascicularis* | 2 | 2 | 6 | 3 | 15 | 13 | 9 | 17 |
| *C. sabaues* | 2 | 0 | 10 | 1 | 7 | 11 | 8 | 8 |
| *P. anubis* | 2 | 3 | 9 | 4 | 18 | 12 | 7 | 13 |
| HUMANOIDES |  |  |  |  |  |  |  |  |
| *N. leucogenys* | 1 | 2 | 6 | 2 | 4 | 4 | 3 | 6 |
| *P. abelii* | 3 | 3 | 9 | 6 | 18 | 13 | 10 | 10 |
| *G. gorilla* | 1 | 2 | 9 | 3 | 6 | 6 | 6 | 11 |
| *P. paniscus* | 1 | 1 | 7 | 3 | 2 | 3 | 5 | 8 |
| *P. troglodytes* | 1 | 2 | 8 | 7 | 8 | 6 | 8 | 12 |
| *H. sapiens* | 1 | 2 | 10 | 5 | 3 | 5 | 7 | 11 |

**Table 3:**
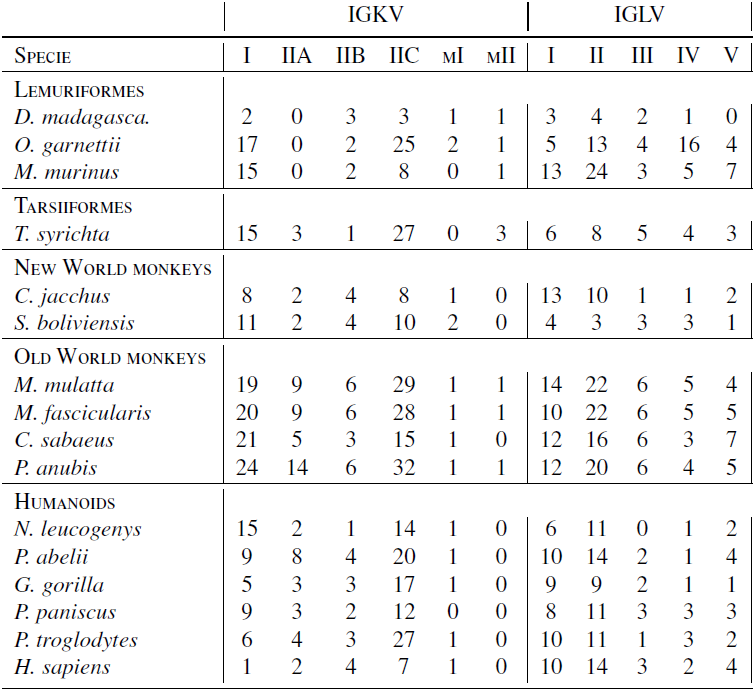
Distribution of V exons from IGKV and IGLV across clades and species.

|  | IGKV |  |  |  |  |  | IGLV |  |  |  |  |
| --- | --- | --- | --- | --- | --- | --- | --- | --- | --- | --- | --- |
| SPECIE | I | IIA | IIB | IIC | MI | MII | I | II | III | IV | V |
| LEMURIFORMES |  |  |  |  |  |  |  |  |  |  |  |
| *D. madagasca.* | 2 | 0 | 3 | 3 | 1 | 1 | 3 | 4 | 2 | 1 | 0 |
| *O. garnettii* | 17 | 0 | 2 | 25 | 2 | 1 | 5 | 13 | 4 | 16 | 4 |
| *M. murinus* | 15 | 0 | 2 | 8 | 0 | 1 | 13 | 24 | 3 | 5 | 7 |
| TARSIIFORMES |  |  |  |  |  |  |  |  |  |  |  |
| *T. syrichta* | 15 | 3 | 1 | 27 | 0 | 3 | 6 | 8 | 5 | 4 | 3 |
| NEW WORLD MONKEYS |  |  |  |  |  |  |  |  |  |  |  |
| *C. jacchus* | 8 | 2 | 4 | 8 | 1 | 0 | 13 | 10 | 1 | 1 | 2 |
| *S. boliviensis* | 11 | 2 | 4 | 10 | 2 | 0 | 4 | 3 | 3 | 3 | 1 |
| OLD WORLD MONKEYS |  |  |  |  |  |  |  |  |  |  |  |
| *M. mulatta* | 19 | 9 | 6 | 29 | 1 | 1 | 14 | 22 | 6 | 5 | 4 |
| *M. fascicularis* | 20 | 9 | 6 | 28 | 1 | 1 | 10 | 22 | 6 | 5 | 5 |
| *C. sabaesus* | 21 | 5 | 3 | 15 | 1 | 0 | 12 | 16 | 6 | 3 | 7 |
| *P. anubis* | 24 | 14 | 6 | 32 | 1 | 1 | 12 | 20 | 6 | 4 | 5 |
| HUMANOIDES |  |  |  |  |  |  |  |  |  |  |  |
| *N. leucogenys* | 15 | 2 | 1 | 14 | 1 | 0 | 6 | 11 | 0 | 1 | 2 |
| *P. abelii* | 9 | 8 | 4 | 20 | 1 | 0 | 10 | 14 | 2 | 1 | 4 |
| *G. gorilla* | 5 | 3 | 3 | 17 | 1 | 0 | 9 | 9 | 2 | 1 | 1 |
| *P. paniscus* | 9 | 3 | 2 | 12 | 0 | 0 | 8 | 11 | 3 | 3 | 3 |
| *P. troglodytes* | 6 | 4 | 3 | 27 | 1 | 0 | 10 | 11 | 1 | 3 | 2 |
| *H. sapiens* | 1 | 2 | 4 | 7 | 1 | 0 | 10 | 14 | 3 | 2 | 4 |

From each of the clades, we determined the 90% consensus sequences. These sequences are represented in the lower part of Figure 2, showing sequence conservation in the framework regions and the existence of motifs that can be used to define separate clades in these regions. As expected, the least conserved regions correspond to those of the CDRs.

We found similar results in the IG light chain V genes. In particular, we found that across the primate species, there was wide variability in number IGKV genes (coding for the κ, or IGK, chains), ranging between 10 and 66 V genes. With respect to the number of IGLV (coding for the λ, or IGL, chain), we found a similar variability, between 10 and 51 V genes. Also, some variation exists between the number ratios IGKV/IGLV within each of the primate species studied. These ratios are of interest, because it is well established that in humans and mice, there is more IGK than IGL both in serum as well as with respect to the number of genes (ie., in humans, there are 46 IGKV genes and 33 IGLV in humans, while in serum there is approximately 70% IGK chain antibodies as compared to 30% IGL antibody chains). In all primate species, we found a larger number of IGKV compared to IGLV genes, with only two exception, in prosimians, in which the ratio is unity, and in the case of the *M. murinus* for which the ratio is inverted with respect to other primates.

From the 16 primate species in our study, we obtained a total of 629 IGKV exon sequences. The resulting phylogenetic tree from the corresponding AA translated sequences indicates the existence of two large (or principal) clades and two smaller clades having a lower number of sequences (Clade I-210 sequences- and II-419 sequences-seen in Figure 3). The two principal clades have representative sequences from all the primates studied, except for prosimians, which do not have sequences in clade IIA.

**Figure 3:**
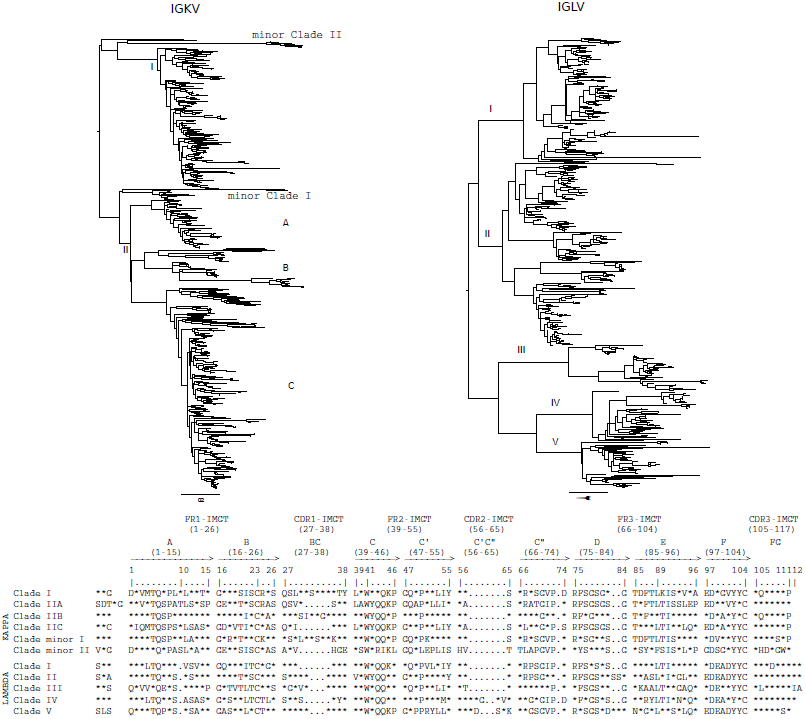
The phylogenetic trees of the AA translated sequences from (IGKV-left- and IGLV-right-) exons from 16 primates species. V exon sequences are obtained from whole genome shotgun (WGS) datasets using the Vgenextractor algorithm. Alignment of the amino acid sequences was performed with clustalO, tree construction with FastTree using the WAG matrix, and visualization with Figtree. The root of the major clades are marked with Roman numerals. Significant subclades in the tree are identified to the right. In the bottom part of the figure, the consensus sequences of each clade are given. The amino acids that are found in more than 90 % of the sequences are marked by their letter, while the variable regions are represented by an asterisk (“*”).

We obtained a total of 522 IGLV exon sequences from the 16 primate species studied. These sequences group into five principal clades, each having representatives from all species. This clade structure may be significant, since it corresponds to the five clades in IGLV locus we described in a previous publication (Olivieri et al., 2014). Indeed, this structural conservation seen in the IGLV clades may have functional significance, because it is also maintained in distant reptiles species.

From sequence alignments within each clade, we deduced motifs from the 90% consensus sequences (those sequences whose AA positions possess 90% similarity) given in Figure 3 (bottom). The AA in the sequences are present in nearly all sequences, while the “*” represents sites of variability. Similar to what we showed for the IGHV exons, the clades are defined by sequence motifs present in the frameworks FR1 and FR3. The sequences in the FR2 region from different clades are similar, while variability can be detected in regions that contain the CDRs.

### 3.2 The TR V genes

We used our gene calling algorithm, *Vgenextractor*, to obtain the TRV exon sequences and study the AA sequences of the V exons from the TRA and TRB loci in 16 different primate species. In particular, we obtained 670 TRAV exon sequences and found that the number of TRAV exons ranges between 30 (in *C. jacchus*) and 69 (in *O. garnettii*). From a phylogenetic study of the AA sequences of the TRAV exons, six major clades can be identified. Also, each of these clades have several subclades. From a detailed examination of these subclades, at least one sequence from each species is found to be common, indicating that these are clades of orthologous genes. Figure 3 shows phylogenetic tree of the AA translated sequences from the TRAV exon sequences, showing a natural grouping into 35 subclades. Most of these subclades have sequences for at least twelve species. In the figure, we zoomed in on specific subclades to expose the taxa to which the constituent sequences originate.

Table 5 lists the V exon sequence distribution for each species per clade. In general, each species is represented within each of the 35 clades with one or two genes. There are clades where the V exon sequences of some species are absent, however this could be because we did not detect the V exon with our gene calling algorithm. In previous publications, we show that our algorithm detects approximately 95% of V exon sequences (Olivieri et al., 2013).

Similar to the method we used in the IG loci, we determined the 90% consensus sequences for each of the 35 clades for V exons in the TRA locus. Clear differences can be seen between sequences from different clades and conservation exist within the same clade. Table 4) shows the variability that exists within each of the V exon sequence regions (FR1, CDR1, FR2, CDR2 and FR3). Within each of these five regions, we determined the fraction of the number of locations having variability (with positions shown as ‘*’) to the total number of positions including the amino acid conserved sites. As expected, the CDR regions are those that exhibit the most AA site variability, however CDR2 is less variable than the CDR1, suggesting an underlying conservation processes. When compared against the framework regions, the CDR2 region is slightly more variable.

**Table 4:** Percentage of AA sites along the translated V exon primate sequences, derived from the alignments provided in Figures 4 and 6.

| Locus | FR1 | CDR1 | FR2 | CDR2 | FR3 |
| --- | --- | --- | --- | --- | --- |
| TRAV | 32% | 54% | 30% | 38% | 30% |
| TRBV | 34% | 43% | 27% | 59% | 31% |
| IGHV | 24% | 54% | 31% | 68% | 29% |
| IGKV | 44% | 56% | 35% | 50% | 32% |
| IGLV | 50% | 85% | 51% | 83% | 39% |

**Table 5:** Number of TRAV exons present in each clade by specie in the phylogenetic tree defined in Figure 4.

| CLADE | *D. madagasc.* | *O. garnettii* | *M. murinus* | *T. syrichta* | *C. jacchus* | *S. boliviensis* | *M. mulatta* | *M. fascicularis* | *C. sabaesus* | *P. anubis* | *N. leucogenys* | *P. abelii* | *G. gorilla* | *P. paniscus* | *P. troglodytes* | *H. sapiens* |
| --- | --- | --- | --- | --- | --- | --- | --- | --- | --- | --- | --- | --- | --- | --- | --- | --- |
| 1 | 0 | 0 | 0 | 0 | 1 | 1 | 0 | 0 | 0 | 0 | 0 | 1 | 0 | 1 | 1 | 1 |
| 2 | 1 | 2 | 1 | 1 | 1 | 1 | 1 | 1 | 1 | 1 | 1 | 1 | 0 | 0 | 0 | 0 |
| 3 | 1 | 1 | 1 | 1 | 1 | 1 | 1 | 1 | 1 | 0 | 1 | 1 | 1 | 0 | 1 | 1 |
| 4 | 3 | 4 | 1 | 0 | 0 | 0 | 1 | 1 | 1 | 1 | 1 | 1 | 1 | 1 | 1 | 1 |
| 5 | 1 | 2 | 0 | 1 | 1 | 1 | 1 | 1 | 1 | 1 | 1 | 1 | 1 | 1 | 1 | 1 |
| 6 | 0 | 0 | 0 | 0 | 1 | 2 | 1 | 1 | 1 | 1 | 1 | 3 | 1 | 1 | 2 | 2 |
| 7 | 2 | 1 | 1 | 2 | 0 | 3 | 2 | 3 | 2 | 3 | 3 | 2 | 2 | 2 | 2 | 3 |
| 8 | 4 | 8 | 5 | 4 | 2 | 3 | 2 | 2 | 1 | 2 | 3 | 3 | 4 | 3 | 4 | 3 |
| 9 | 1 | 2 | 5 | 1 | 1 | 1 | 1 | 1 | 1 | 1 | 1 | 1 | 1 | 1 | 1 | 1 |
| 10 | 1 | 0 | 2 | 1 | 1 | 1 | 1 | 1 | 1 | 1 | 1 | 1 | 1 | 1 | 1 | 1 |
| 11 | 1 | 1 | 1 | 1 | 1 | 1 | 2 | 2 | 1 | 1 | 1 | 2 | 1 | 0 | 2 | 1 |
| 12 | 0 | 1 | 0 | 0 | 1 | 1 | 0 | 0 | 0 | 0 | 1 | 0 | 0 | 0 | 0 | 0 |
| 13 | 2 | 2 | 0 | 2 | 1 | 1 | 1 | 1 | 1 | 1 | 1 | 1 | 1 | 1 | 1 | 1 |
| 14 | 2 | 5 | 1 | 2 | 0 | 0 | 1 | 1 | 1 | 1 | 1 | 1 | 1 | 1 | 2 | 1 |
| 15 | 1 | 15 | 0 | 1 | 1 | 0 | 2 | 2 | 2 | 2 | 2 | 2 | 2 | 2 | 2 | 2 |
| 16 | 0 | 2 | 0 | 1 | 1 | 2 | 2 | 2 | 1 | 2 | 1 | 3 | 2 | 1 | 2 | 2 |
| 17 | 1 | 1 | 0 | 2 | 1 | 1 | 1 | 1 | 3 | 0 | 2 | 2 | 1 | 1 | 1 | 1 |
| 18 | 1 | 1 | 1 | 1 | 0 | 1 | 1 | 1 | 1 | 1 | 1 | 1 | 1 | 1 | 1 | 1 |
| 19 | 0 | 1 | 0 | 0 | 1 | 1 | 1 | 1 | 0 | 1 | 1 | 1 | 1 | 1 | 1 | 1 |
| 20 | 1 | 0 | 2 | 2 | 0 | 0 | 1 | 1 | 1 | 1 | 1 | 1 | 1 | 1 | 1 | 1 |
| 21 | 3 | 5 | 3 | 2 | 3 | 3 | 2 | 3 | 2 | 3 | 3 | 2 | 3 | 3 | 3 | 3 |
| 22 | 1 | 1 | 1 | 1 | 1 | 0 | 1 | 1 | 1 | 1 | 1 | 1 | 1 | 1 | 1 | 1 |
| 23 | 1 | 1 | 1 | 1 | 0 | 0 | 1 | 1 | 1 | 1 | 1 | 1 | 1 | 1 | 2 | 1 |
| 24 | 1 | 0 | 1 | 1 | 1 | 1 | 1 | 1 | 1 | 1 | 1 | 1 | 1 | 1 | 1 | 1 |
| 25 | 0 | 0 | 0 | 1 | 0 | 0 | 1 | 1 | 0 | 1 | 1 | 1 | 1 | 1 | 1 | 0 |
| 26 | 1 | 1 | 1 | 1 | 1 | 1 | 1 | 1 | 1 | 1 | 1 | 0 | 1 | 1 | 1 | 1 |
| 27 | 0 | 0 | 0 | 0 | 1 | 1 | 1 | 1 | 1 | 1 | 1 | 1 | 1 | 1 | 1 | 1 |
| 28 | 0 | 0 | 0 | 1 | 1 | 1 | 1 | 1 | 1 | 1 | 0 | 0 | 1 | 0 | 2 | 1 |
| 29 | 1 | 1 | 3 | 1 | 0 | 0 | 2 | 2 | 1 | 1 | 1 | 1 | 1 | 0 | 3 | 1 |
| 30 | 1 | 0 | 0 | 0 | 1 | 1 | 2 | 2 | 1 | 1 | 1 | 0 | 1 | 1 | 2 | 1 |
| 31 | 1 | 1 | 1 | 1 | 1 | 1 | 1 | 1 | 1 | 1 | 1 | 1 | 1 | 1 | 2 | 1 |
| 32 | 0 | 0 | 0 | 2 | 1 | 1 | 1 | 1 | 1 | 1 | 2 | 1 | 2 | 2 | 2 | 2 |
| 33 | 1 | 0 | 0 | 1 | 0 | 1 | 1 | 1 | 1 | 1 | 1 | 1 | 1 | 1 | 1 | 1 |
| 34 | 1 | 0 | 0 | 0 | 1 | 1 | 1 | 1 | 0 | 1 | 1 | 1 | 1 | 1 | 1 | 1 |
| 35 | 4 | 7 | 4 | 2 | 1 | 1 | 2 | 1 | 1 | 1 | 1 | 1 | 1 | 1 | 1 | 1 |

**Figure 4:**
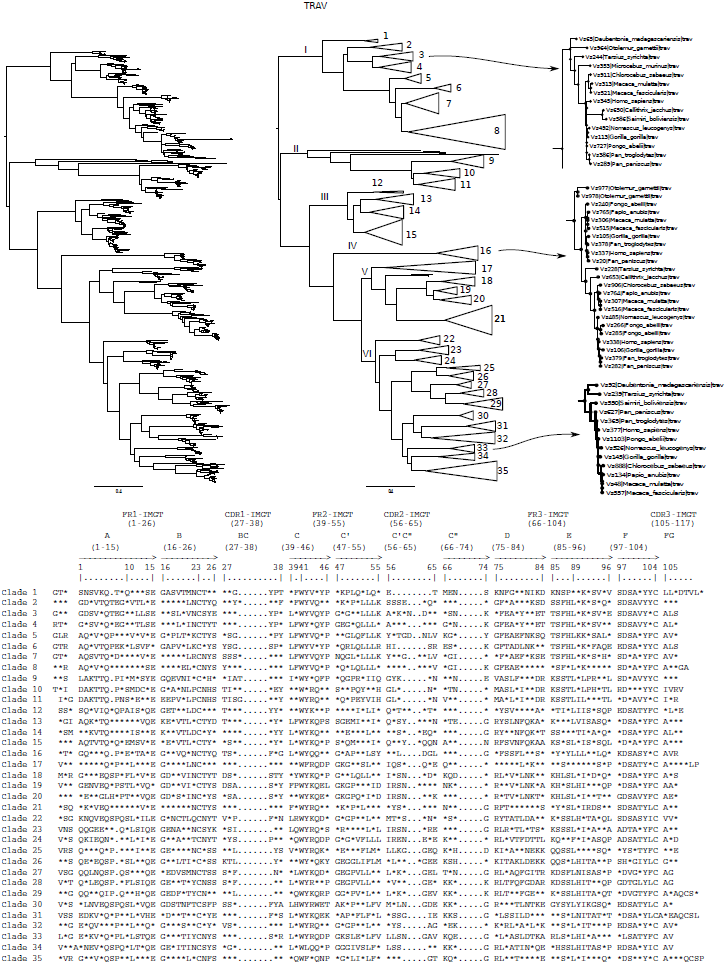
The phylogenetic trees of the AA translated sequences from TRAV exons from 16 primates species. V exon sequences are obtained from whole genome shotgun (WGS) datasets using the Vgenextractor algorithm. Alignment of the amino acid sequences was performed with clustalO, tree construction with FastTree using the WAG matrix, and visualization with Figtree. The original tree with all V exon sequences are shown to the left. Clades are collapsed in the center tree to better illustrate the sequence similarities. Representative subclades, 8 and 23, are shown to the right to illustrate the distribution of constituent primate taxa. At the bottom part of the figure, the consensus sequences of each clade are shown, where the amino acids that are found in more than 90% of the sequences are marked by their letter, while the variable regions are represented by an asterisk (“*”)

**Figure 5:**
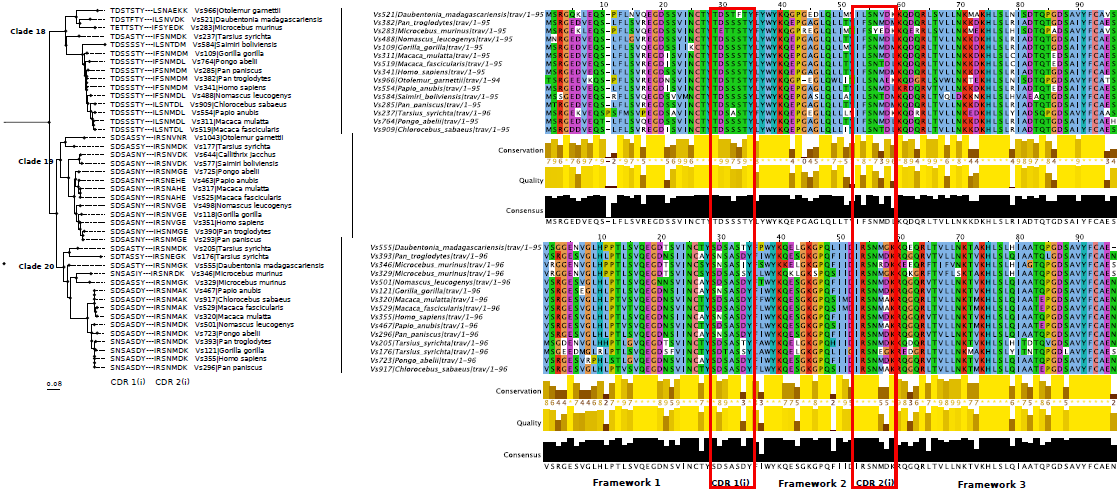
Detailed representation of the clades 18, 19 and 20 formed from the alignment of the AA sequences of TRAV exon sequences of primates. The phylogenetic tree (left) show the CDR1(i) and CDR2 (i) sequences in order to demonstrate the similarity of these sequences between members of the same clade. Sequence alignment (right) is shown for clade-18 and clade-20. The regions that are marked have been defined by our analysis software, Trozos, (see material and methods).

For TRBV exon sequences, we obtained similar results with respect to clade grouping as we found for the TRA locus. In particular, we obtained 696 TRBV exon sequences and we can deduce 25 clades from a tree analysis. As in the previous cases discussed, we found that all clades contain V sequences from the majority of primate species. Table 6 shows the distribution of AA translated TRBV exon sequences across the different subclades for species. As can be seen, each specie contributes one or more sequences per clade. Figure 6 shows the phylogenetic tree of all the AA translated TRBV exon sequences, together with the alignment within each clade to obtain the 90 % consensus sequences. As before, we studied the variability between the canonical IMGT defined regions. The results are shown in Table 4, where it can be seen that the greatest variability is detected within the CDRs. Also, in amongst the TRAV genes, the sequences of CDR1 have lower variability than those of CDR2.

**Figure 6:**
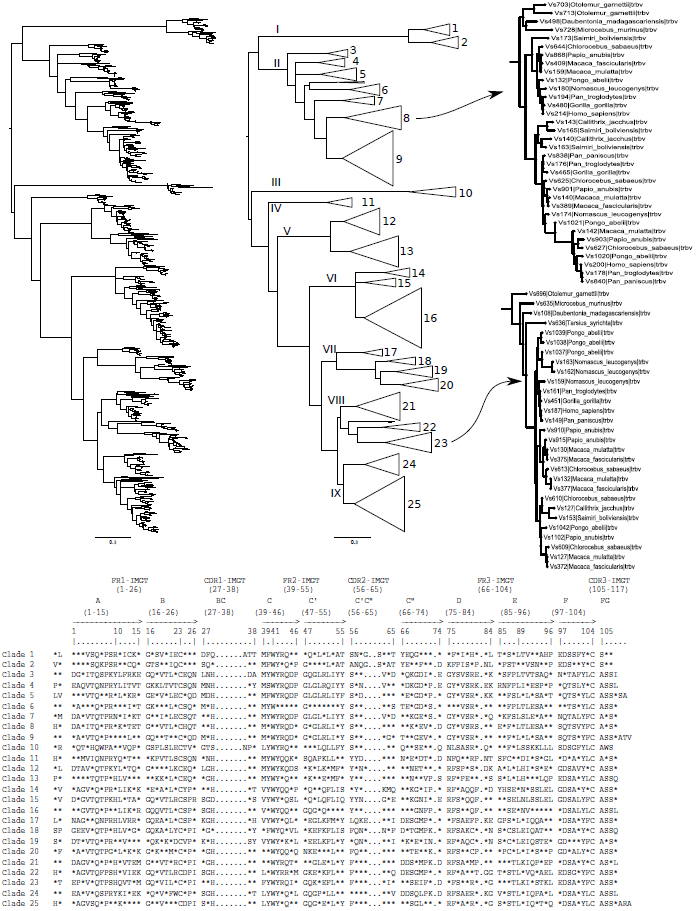
The phylogenetic trees of the AA translated sequences from TRAV exons from 16 primates species. V exon sequences are obtained from whole genome shotgun (WGS) datasets using the Vgenextractor algorithm. Alignment of the amino acid sequences was performed with clustalO, tree construction with FastTree using the WAG matrix, and visualization with Figtree. The original tree with all V exon sequences are shown to the left. Clades are collapsed in the center tree to better illustrate the sequence similarities. Representative subclades, 3, 16, and 33, are shown to the right to illustrate the distribution of constituent primate taxa. At the bottom part of the figure, the consensus sequences of each clade are shown, where the amino acids that are found in more than 90% of the sequences are marked by their letter, while the variable regions are represented by an asterisk (”*”).

**Table 6:** Number of TRBV genes present in each primate species in the phylogenetic clades defined in Figure 6.

| CLADE | *D. madagasc.* | *O. garnettii* | *M. murinus* | *T. syrichta* | *C. jacchus* | *S. boliviensis* | *M. mulatta* | *M. fascicularis* | *C. sabaeus* | *P. anubis* | *N. leucogenys* | *P. abelii* | *G. gorilla* | *P. paniscus* | *P. troglodytes* | *H. sapiens* |
| --- | --- | --- | --- | --- | --- | --- | --- | --- | --- | --- | --- | --- | --- | --- | --- | --- |
| 1 | 1 | 1 | 0 | 0 | 1 | 1 | 1 | 1 | 1 | 1 | 1 | 1 | 1 | 2 | 2 | 2 |
| 2 | 1 | 1 | 1 | 1 | 1 | 1 | 1 | 1 | 1 | 1 | 1 | 1 | 1 | 2 | 2 | 1 |
| 3 | 0 | 0 | 0 | 1 | 1 | 1 | 1 | 1 | 1 | 1 | 0 | 1 | 1 | 1 | 1 | 1 |
| 4 | 0 | 1 | 0 | 0 | 1 | 1 | 1 | 1 | 1 | 1 | 1 | 1 | 1 | 1 | 1 | 1 |
| 5 | 1 | 1 | 1 | 1 | 1 | 1 | 3 | 1 | 1 | 1 | 1 | 2 | 1 | 1 | 1 | 1 |
| 6 | 1 | 0 | 0 | 0 | 1 | 2 | 1 | 1 | 1 | 1 | 1 | 3 | 2 | 2 | 2 | 2 |
| 7 | 0 | 1 | 1 | 1 | 1 | 0 | 1 | 1 | 1 | 1 | 0 | 1 | 1 | 1 | 1 | 1 |
| 8 | 1 | 2 | 1 | 0 | 2 | 3 | 2 | 2 | 3 | 3 | 2 | 3 | 2 | 2 | 3 | 2 |
| 9 | 0 | 4 | 2 | 0 | 3 | 3 | 3 | 5 | 6 | 7 | 4 | 8 | 6 | 8 | 8 | 6 |
| 10 | 1 | 1 | 1 | 0 | 1 | 1 | 1 | 1 | 1 | 1 | 1 | 1 | 1 | 1 | 1 | 1 |
| 11 | 1 | 1 | 1 | 0 | 1 | 1 | 1 | 1 | 1 | 1 | 1 | 1 | 1 | 0 | 1 | 1 |
| 12 | 0 | 2 | 4 | 1 | 2 | 1 | 2 | 4 | 3 | 4 | 3 | 3 | 2 | 2 | 2 | 1 |
| 13 | 1 | 3 | 4 | 0 | 4 | 1 | 4 | 3 | 3 | 3 | 4 | 4 | 2 | 3 | 3 | 3 |
| 14 | 1 | 1 | 0 | 1 | 1 | 1 | 1 | 1 | 1 | 1 | 1 | 0 | 1 | 1 | 1 | 1 |
| 15 | 0 | 0 | 0 | 1 | 1 | 1 | 1 | 1 | 1 | 1 | 1 | 1 | 1 | 0 | 1 | 1 |
| 16 | 1 | 2 | 5 | 0 | 5 | 4 | 5 | 7 | 8 | 8 | 4 | 7 | 6 | 7 | 7 | 6 |
| 17 | 1 | 1 | 0 | 0 | 1 | 1 | 1 | 1 | 0 | 0 | 0 | 1 | 1 | 1 | 1 | 1 |
| 18 | 1 | 0 | 0 | 0 | 1 | 1 | 1 | 1 | 1 | 1 | 1 | 1 | 0 | 1 | 1 | 1 |
| 19 | 1 | 1 | 1 | 0 | 1 | 1 | 1 | 1 | 1 | 1 | 0 | 3 | 2 | 1 | 1 | 1 |
| 20 | 1 | 1 | 1 | 0 | 1 | 1 | 1 | 1 | 1 | 1 | 1 | 1 | 2 | 2 | 2 | 2 |
| 21 | 1 | 2 | 1 | 1 | 4 | 3 | 4 | 3 | 3 | 3 | 1 | 5 | 3 | 2 | 2 | 3 |
| 22 | 0 | 0 | 1 | 0 | 1 | 1 | 1 | 1 | 1 | 1 | 1 | 1 | 1 | 1 | 1 | 1 |
| 23 | 1 | 1 | 1 | 1 | 1 | 1 | 1 | 3 | 3 | 3 | 3 | 4 | 1 | 1 | 1 | 1 |
| 24 | 0 | 1 | 1 | 2 | 2 | 0 | 2 | 3 | 3 | 3 | 1 | 2 | 2 | 2 | 3 | 3 |
| 25 | 1 | 3 | 3 | 1 | 4 | 4 | 4 | 9 | 6 | 9 | 2 | 5 | 7 | 6 | 6 | 5 |

The phylogenetic results from the TRBV and TRAV loci show that V exons sequences exist that are maintained throughout evolution across primate species, since each species contributes one such gene to the subclades of the tree. To confirm these results and to study which parts of the sequences are involved in the process of selection, we studied independently the framework and CDR sequences separately from each V exon sequence. To separate the four separate the FR1, FR2, FR3, and the CDRs from the AA translated V exon sequences, we developed a python utility program, called Trozos.py. Three sequences correspond to framework regions FR1, FR2 and FR3, while the fourth sequences is constructed by combining the two sequences from CDR1 and CDR2.

Once the sequence fragments FR1, FR2, FR3, and the CDRs were separated, we studied whether each V exon of the TRV is unique to each species and whether it has an ortholog in other species (as suggested by the results of the phylogenetic trees). Figure 7 shows the particular case of a randomly selected V exon sequence for illustration (Vs367 of *H. sapiens* for the TRAV and Vs168 of *M. mulatta* from the TRBV locus; sequences can be obtained from http://vgenerepertoire.org). For example, in the case of the TRAV sequence shown (ie., Vs367 *Homo sapiens*), each of the 4 segments (CDRS, FR1, FR2 and FR3) differ significantly from other TRAV sequences within the same species, appearing as an outlier in the boxplot of Figure 7. In 12 primate species, we found one or two sequences which are similar, indicating that that they are orthologs. This phenomenon occurs in each of the segments, indicating the uniqueness of each V gene. We repeated the same experiment for the TRBV genes (ie., the Vs168 exon sequence of *Macaca mulatta*, shown in Figure 7 (bottom)). The results are similar to those of the TRAV, however unique V exon sequences were not found in the FR2 regions.

**Figure 7:**
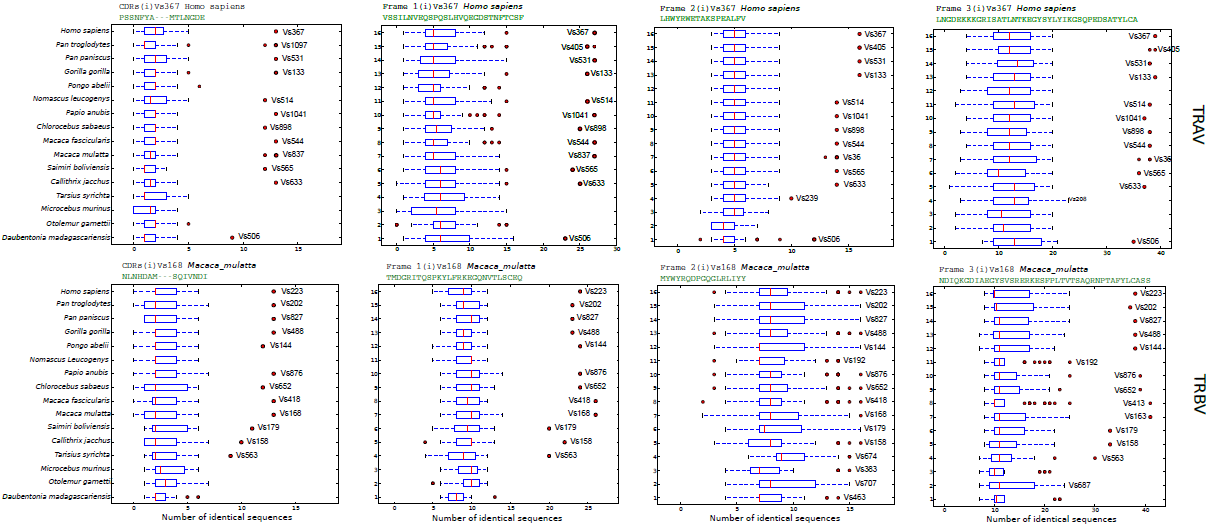
Participation of each of the canonical segments, FR1, FR2, FR3, CDR1, and CDR2, from the amino acid translated V exon sequences of the uniquely identical genes. From these sequences, we obtained the framework regions (1, 2 and 3) and other sequences created artificially with the two CDRs (1 and 2). A concrete case of sequences, originating from a TRAV sequence (top) was compared (identity number) with the rest of the fragments obtained from all TRAV exons from all species. The same experiment was performed with specific sequences of a TRBV (bottom).

The data generated from the phylogenetic tree suggests frequent changes in the gene loci of IG as well as a reduced permissiveness in the genes of the TR chains. The theory of birth and death of genes has been postulated as a mechanism that directs the evolutionary processes of these genes. In Figure 8 we studied this hypothesis by quantifying sequence similarities higher than 90 % in segments over 3000 bases in the IGHV, TRAV and TRBV loci between the orangutan, human and macaque species. The results show that In the IGHV locus, there are more tracks and cross linking as compared to the TR loci. Also, comparing species uncovers relationships in the IGHV locus over evolutionary time. For example, the number of IGHV tracks and crossovers is higher between human and macaque (more distantly related species) than between human and orangutan (more evolutionarily close species). In the TRAV and TRBV loci, the tracks are approximately parallel, indicating that in these loci, less duplication/deletion processes took place between speciation events, contrasting what can be observed in the IGHV locus.

**Figure 8:**
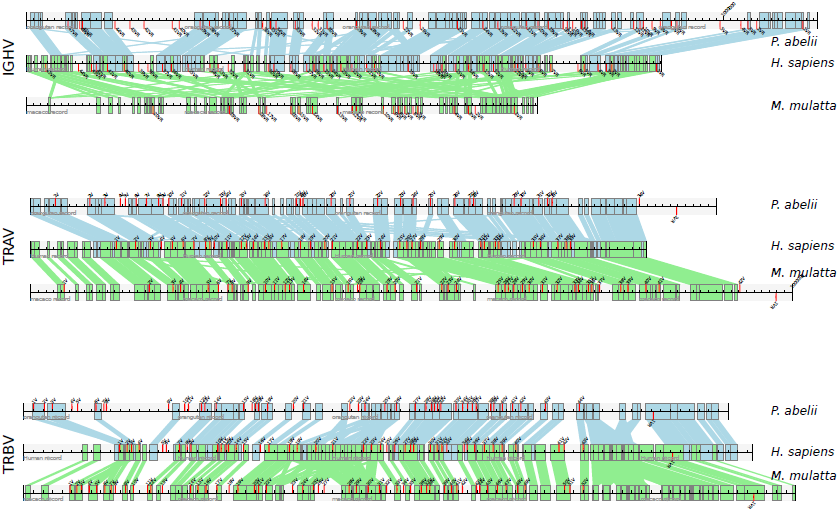
Identical Sequence within the IGHV, TRAV and TRBV loci from the three species of primates, *H. sapiens*, *P. abelii* and *M. mulatta*. For each species, the V exon sequences were extracted from genomic segments available at the Ensemble repository www.ensembl.org. For our analysis pipeline, we used Galaxy (http://galaxy.wur.nl). Sequences for the tracks were obtained in the following way: we performed a BLASTN against the orangutan and macaque sequences (ie., the db), with the query consisting of human sequence. We selected sequence identities > 60% and an alignment length > 3000 bases and the figure was made with a custom python script. The locations of the V exons, marked in red, were obtained with Vgenextractor (http://vgenerepertoire.org/).

## 4 Discussion

In previous studies (Olivieri et al., 2013, 2014), we showed data indicating a different evolutionary process between the V genes of IG and TR. In IG, the processes of birth and death are quite evident. We also highlighted the grouping of sequences into established IMGT clans and proposed new clades. For the species studied in this work, we describe the clustering of the IG light chain sequences into major clades. While these groupings are not as obvious as the three clans of the IGH chains, we can establish these light chain clades with certainty due to the large number of sequences, supporting their existence. The grouping of the IGL chains into five clades is of particular interest, since these clades originated prior to the diversification of mammals and reptiles and interestingly both have remained in evolutionary lines for over 300 million years, suggesting a functional significance of each clade which is still unknown.

All loci containing V exons are very similar. Despite this wide similarity, there are stark evolutionary differences amongst the IG and TR loci. The IG loci exhibit a more pronounced rate of change as than the TR loci. This is seen by observing sequences between species of primates where there is greater sequence conservation in the TCR loci. Besides the evidence left as relics in genomic sequences, frequent duplications of IG genes generate recent clades with multiple members. Evolution provides a defense mechanism of an organism for rapid adaptation of IG chains to a rapidly changing external infectious environment.

In the TRA and TRB loci, there is a conservation of V exons and a low duplication permissiveness. In particular, we found a conservation of 35 TRAV exon sequences and 25 TRBV exon sequences. Nonetheless, in some species, we did not detect any conserved V exons. This may be a methodological error (Vgenextractor only detects 95 % of the V exons from WGS data sets) or the system may be slightly redundant, permitting some V exon loss without compromising the survival of the individual. Similarly, in the TR loci we detected duplication events but never observed multiple duplications, such as those in the IG loci.

The uniqueness of each gene in the TR loci is of particular interest. The number of V genes from these loci is not arbitrary. The fact that a large repertoire variation can be generated by the process of VDJ recombination and somatic mutation has given rise to the assumption that a few V genes should be sufficient for somatic diversification. Previous publications (Suárez et al., 2006) suggest that few V regions can generate nearly complete repertoires. The results expressed in this work indicate that the genomic diversity of the V genes in the TR loci should have a functional basis maintained throughout evolution.

Our results also provide new insights into the evolution of CDR and framework regions. From an evolutionary point of view, the CDRs are sequence segments that should be permissive to mutations, while changes in the framework regions should be less permissive since they provide a well defined structure. In general, when the AA sequences deduced of the V exons are aligned, the CDR regions are grouped in regions called hypervariable regions. However, when each clade is studied independently, the framework regions have a variability similar to that found in the CDR regions (Figure 5) especially in the CDR2 of the TRAV locus and the CDR1 of the TRBV locus. These results show that sequences of this CDRs are maintained in evolution and that there is not a greater permissiveness to mutations than in framework regions. The hypervariability found in the alignment of sequences of one specie is due to the presence of different CDRs within each V exon, but there exists an evolutionary ortholog maintained in other primate species. In Table 5, a column with the consensus sequences of the CDRs (i) are shown. This data indicates that the sequences of each TRAV exon may be positively selected with a specific, non-redundant function.

Why have these genes been maintained in the TRV loci?. A probable explanation is that this maintenance is due to a co-evolution with interacting molecules, such as MHC, that provide a natural evolutionary pressure. This same evolutionary pressure may also condition the pairing of the TRA and TRB regions. Therefore, the evolution of each V region must be constrained by modifications that equally occur in the MHC molecules as well as other changes in V region pairing. These same mechanisms do not occur in the IG loci, since antigen recognition by antibodies is not restricted by MHC molecules, making it likely the greater permissibility towards evolutionary modifications.

Why are there 25 TRBV and 35 TRAV?. A possible explanation could be that a minimum number of genes are required to form TRA/TRB pairs needed for T lymphocytes to recognize antigen presented by the large structural variations of MHC class I and class II molecules. Indeed, it is known that MHC can have multiple forms, particularly class II molecules. If this hypothesis were true, we would expect to find specific pairings of TRA/TRB for putative MHC molecules. Also, we would expect to find evidence of the association between V exons and the presence or absence of MHC genes in evolutive studies.

A plausible explanation for the result we presented are that the MHC genes that coexist with the TRV genes must act as evolutionary guides. In this scenario, the capacity for the TR to recognize the MHC should be coded directly within the germline, while the antigen recognition of the TR-MHC complex is a consequence of random somatic variations in the individual (VDJ re-arrangements and somatic mutation). Studies suggest that recognition of MHC is mediated by the CDR1 and CDR2 which are within the V exon, while the antigenic component is recognized by the CDR3 (encoded by D and J exons) (Marrack et al., 2008; Deng et al., 2012). Our data is consistent with this description and that MHC recognition system must be encoded in the genome. This would explain the coevolution of both molecules. It is logical that the processes of somatic variability are directed towards antigen recognition and have a stochastic quality. If this were the case, the MHC recognition structures would be limited in order to accompany evolutionary allowed changes. Our work points to the fact that these constraining structures are the FR and CDR amino acid sequences generated from the V exons.

## 6. Supplementary Online Material

**Table 1:**
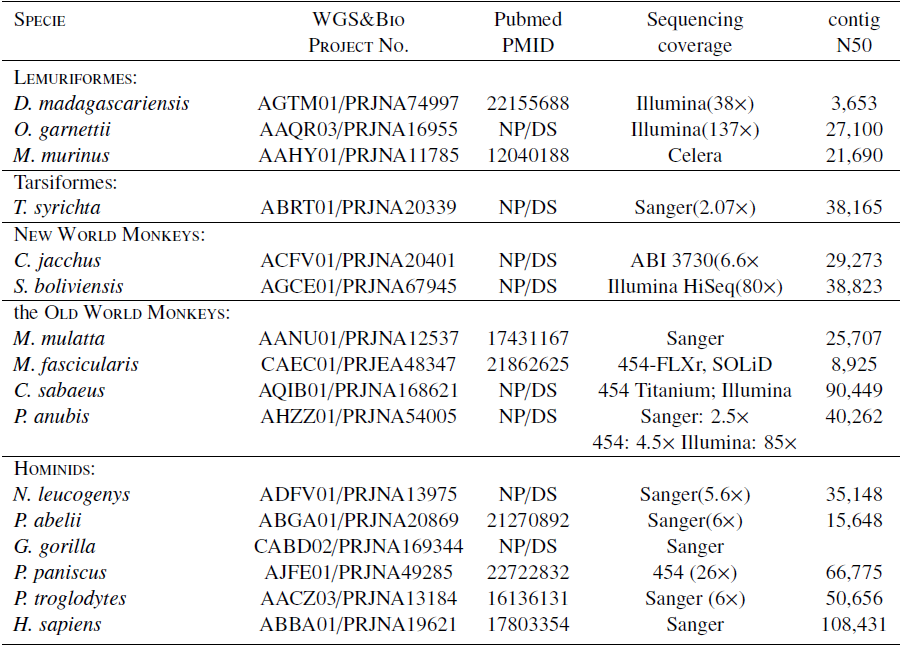
WGS Data for the 16 primates included in this study.NP/DS indicates no publication, direct submission.

